# Spatial resource heterogeneity increases diversity and evolutionary potential

**DOI:** 10.1101/148973

**Authors:** Emily L. Dolson, Samuel G. Pérez, Randal S. Olson, Charles Ofria

## Abstract

Spatial heterogeneity is believed to be an evolutionary driver of biodiversity. Variability in the distribution of resource patches can allow an environment to support a wider variety of phenotypes for selection to act upon at the ecosystem level, which may lead to more species. However, the generality of this principle has not been thoroughly tested, as the relevant adaptive dynamics occur on evolutionary timescales. We overcame this challenge by performing experiments on populations of digital organisms in the Avida Digital Evolution Platform, in which we investigated the impact of spatial resource heterogeneity on phenotypic diversity. Since an important benefit of diversity may be increased evolutionary potential, we also tracked the probability of a complex trait evolving in the context of various levels of spatial heterogeneity. We found that spatial entropy and phenotypic diversity have a strong positive correlation and this relationship is consistent across various spatial configurations. Diversity also increases evolutionary potential, but has a much smaller impact than other components of environmental composition. The most important of these components was the mean number of resources present in locations across the environment, likely owing to the importance of building blocks for the evolution of complex features. These results suggest that a general relationship exists between spatial heterogeneity and diversity, beyond the specific ecosystems and timescales in which it has previously been studied. By examining this relationship in the context of phenotypic evolution, we advance a mechanistic understanding of the resulting dynamics. Moreover, our results suggest that the likelihood of evolving various traits can be impacted by the spatial configuration of patches in which these traits are advantageous. These findings have implications for both evolutionary biology and evolutionary computation, as generating and maintaining diversity is critical to all forms of evolution.

## Introduction

A central problem in ecology is understanding the mechanisms that promote species coexistence and richness in natural ecosystems [1, 2]. There is mounting evidence that spatial heterogeneity (the lack of homogeneity across space) is an important driver of biological diversity [3]. A common hypothesis, here termed the positive heterogeneity-diversity hypothesis, states that even seemingly small amounts of spatial heterogeneity promote diversity by creating new spatially-restricted niches for specialist species to inhabit, thereby reducing competition for limited resources [3–5]. This hypothesis is supported by many cases in which spatial heterogeneity clearly influences the dynamics of species coexistence, allowing for greater species diversity than in homogeneous environments [6–10].

Contrary to the positive heterogeneity-diversity hypothesis, some recent studies have found evidence that diversity should peak at intermediate levels of spatial heterogeneity [11–13]. These results are driven by two related factors: 1) the area-heterogeneity tradeoff and 2) decreasing contiguity of any given habitat.

The area-heterogeneity tradeoff is an observation that increasing the number of unique regions within an environment must decrease the average area spanned by each, if total geographic area is held constant [12]. In thinking through the effects of such a decrease, it is important to consider whether these regions represent: A) fundamental niches - the full range in which a species is capable of surviving, or B) preferred habitat - a set of conditions to which a species may be particularly well-optimized to outcompete others. Research suggesting that the area-heterogeneity tradeoff leads to a peaked heterogeneity-diversity relationship has so far only considered heterogeneity in fundamental niches for species of interest [11,12]. Such a scenario is akin to dynamics theorized to occur in harsh climates, in which environmental gradients impose strict constraints on where a species can and cannot survive. The amount of space that is within a species fundamental niche places a cap on its maximum population size. Since smaller populations have an increased risk of stochastic extinction, species whose fundamental niches occupy less area (relative to the ideal amount of area they need) are at greater risk of being lost from the community. These extinctions are believed to drive decreased diversity at high levels of heterogeneity. However this effect should not hold if we instead focus on heterogeneity of preferred habitats. If all species are capable of surviving anywhere in the environment, any stochastic extinction will allow for subsequent range expansion by other species and a corresponding drop in their risk of stochastic extinction. This process leaves an unexploited niche for future speciation. These factors should balance out such that, at worst, diversity remains constant as heterogeneity increases (this hypothesis is supported by the exploration of niche width in [12]). This scenario is more akin to dynamics observed in the tropics, where species realized niches are thought to be constrained primarily by competition. While these two scenarios are both important to understand, and likely represent extrema of a continuum, here we will focus on preferred habitat heterogeneity.

Some metrics of environmental heterogeneity, such as edge-density, define more fragmented environments as more heterogeneous. In a system that defines heterogeneity along only a single environmental gradient, high values of these metrics can be achieved only if the environment is highly fragmented (as in [13]). Similar to the area-heterogeneity tradeoff, this decreased patch contiguity increases the likelihood of stochastic extinctions, reducing population diversity at high levels of heterogeneity. However, when niches are defined along more than one dimension, it is possible to increase heterogeneity without meaningfully increasing fragmentation. Fragmentation is an important issue to study, but it is only one component of heterogeneity. For simplicity, here we will assume that patches of any one resource are large enough to support a stable population, although intersections where multiple patches overlap may not be.

The development of a general theory of the impact of spatial heterogeneity on diversity has been inhibited by the fact that the causal mechanism, populations evolutionarily diversifying into new spatial niches, occurs over a long time scale. As evolution is the origin of biological diversity, a thorough understanding of the effects of spatial heterogeneity on diversity requires consideration of the associated evolutionary dynamics. Evolution occurs on a broad temporal scale, while the relevant ecological dynamics occur on a broad spatial scale. Dealing with either of these scales in a biological experiment is challenging, and their combination is effectively intractable. For this reason, we turn to a computational system to perform *in silico* experiments with digital organisms. In digital evolution, rapid generation times and transparent data allow us to perform large-scale experiments while perfectly controlling resource distributions and measuring organismal phenotypes. Here, we use the Avida Digital Evolution Platform [14]. Avida provides a rich environment for studying evolutionary processes and has been used in the past to investigate phenomena such as the origins of complex features [15], mutational robustness [16], adaptive radiation [17], and evolving ecological networks [18].

Digital organisms in Avida are self-replicating computer programs that can evolve to metabolize limited resources by performing associated mathematical functions. We define the phenotype of a digital organism as the set of resource types that organism can use. Previous studies in Avida have found that introducing negative frequency dependence via resource competition is sufficient to generate and maintain diverse phenotypes through adaptive radiation into multiple niches [17,19]. In this study, we explore the effect of introducing a spatial component to this resource competition. Each organism in Avida occupies one cell in a two-dimensional grid. We create spatial heterogeneity by varying the set of resources available in each grid cell. Specifically, we place circular patches of resources in the environment. Since many patches will overlap, this placement technique creates many new spatial niches. We expect that subsets of the population will adapt to be able to use locally available resources. This process is closely related to ecological speciation [20], but, as speciation is challenging to define for large and complex populations of asexual organisms such as ours, we will focus instead on the diversity of distinct phenotypes. Prior research on speciation in spatially heterogenous environments has demonstrated that some types of heterogeneity can indeed promote evolutionary branching [13]; that study found a peak in diversity at intermediate levels of heterogeneity, which we expect is the result of focusing on heterogeneity along a single environmental gradient.

### Evolutionary potential

Here focus on short-term evolutionary potential, which we define as the probability that a mutated offspring possesses new beneficial adaptations. It should be clear that the diversity of a community will be entangled with the combined evolutionary potential of its constituent populations: On one hand, diversity generally increases evolutionary potential [21, 22], since a community that is more spread out across a genotypic or phenotypic space is more likely to contain an organism that is a short mutational distance away from a given feature. There is also substantial empirical evidence from the field of evolutionary computation that diversity is important to finding globally optimal solutions when using evolution as a problem-solving tool [23–26]. Evolutionary computation, as a context, allows us to sensibly talk about how “useful” diversity is as a whole and which types of diversity are most effective toward achieving the desired goal [26]. This concept can also be translated back to biological evolution in contexts where we are interested in the evolution of a specific feature or set of features (such as increased complexity, or recovery of community traits lost due to a severe disturbance). Thus, understanding how to promote the evolutionary generation of diversity has important practical applications. Conversely, greater evolutionary potential facilitates the evolutionary generation of diversity, as populations that are able to adapt rapidly can take advantage of new niches more quickly.

A previous study in Avida on the relationship between the availability of resources and the resulting phenotypic diversity found that evolutionary potential peaks when all resources are intermediately plentiful. Interestingly, diversity peaks at lower resource levels than evolutionary potential [27]. These results are inconclusive, however, as to whether a direct relationship exists between diversity and evolutionary potential, and highlight the complexity of the relationship between diversity, resource availability, and evolutionary potential. Here, we attempt to clarify the causal relationships between these factors.

### Quantifying environmental heterogeneity

Many metrics have been proposed to quantify spatial heterogeneity, but most fall into two categories: composition and spatial configuration [28]. Composition metrics ignore the relative positions of resources, and instead deal with what types of spatial niches are present in an ecosystem. For example, commonly used composition metrics include *richness* (a count of how many distinct niches exist), *evenness* (do all niches cover the same amount of space?), and composites of the two such as *Shannon entropy*. For the purposes of this study, we use Shannon entropy of the number of cells in which each set of resources is present as a metric of environmental composition. This metric balances both richness and evenness and is a generally accepted metric of spatial heterogeneity [29].

Spatial configuration metrics, on the other hand, deal with arrangements of niches in space and include *patch shape complexity, core area, contrast*, and *isolation*. There is no single metric that can accurately summarize spatial configuration—for any given study it is important to identify which variables are meaningful. Here, we track the extent to which single-resource patches overlap to form multi-resource spatial niches. Because we define an organism’s phenotype as the set of resources that it is able to consume, a spatial niche containing more resource types will be hospitable to a wider variety of phenotypes. Moreover, because of the tendency of simpler functions in Avida to facilitate the evolution of more complex ones, niches with more types of resources will be easier for populations to adapt to. Thus, spatial resource overlap is the primary configuration metric we track here. This overlap is captured by the distribution created by counting the number of resources in each cell of an environment. We will discuss overlap in terms of the mean, variance, skew, and kurtosis of this distribution.

## Results and Discussion

### Niche stability

Our goal is to isolate the effects of heterogeneity from the effects of fragmentation. To do so, we must first determine the minimum patch size that would be required to stably support populations specialized to live in those regions.

Likewise, we need to identify the minimum overlap between two patches to ensure the resulting spatial niche is similarly stable. We measured these threshold sizes empirically by carrying out two series of experiments in which the world contained only two circular resource patches. In the first set of experiments, we tested all possible pairs of resources at three levels of overlap: complete, half-way, or none (Fig 1). For each region, we defined the optimal phenotype as one that performed a set of functions that perfectly matched the available resources.

**Figure 1.**
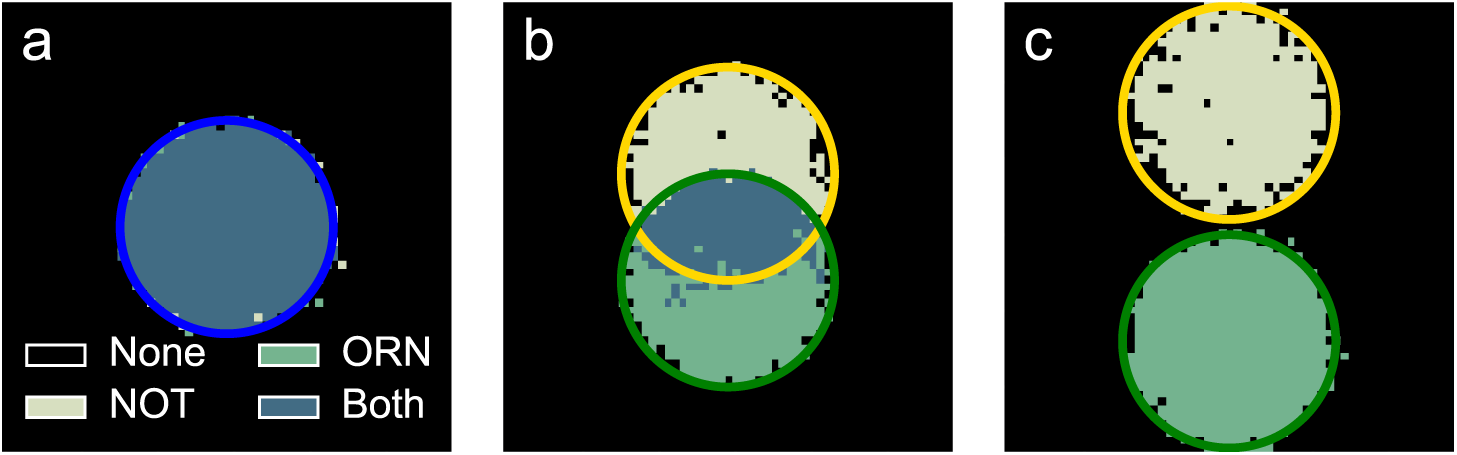
Spatial arrangement of phenotypes at the end of each experiment in worlds containing patches of two different resources, NOT and ORN, each radius 14. Patch locations are indicated by circles - gold represents the NOT resource, green represents the ORN resource, and blue represents both resources overlapping. The color of each cell in the grid indicates the most common phenotype in that cell across the 30 replicates. **A:** Patch centers are 0 cells apart, creating complete overlap. **B:** Patch centers are 14 cells apart, leading to three coexisting phenotypes. **C:** Patch centers are 30 cells apart, resulting in complete patch separation and no phenotype capable of both functions.

The optimal phenotype for the spatial niche at the intersection of the two resource patches evolved consistently when the organisms needed to perform sufficiently simple functions (Fig 1). For example, the optimal phenotype evolved 66% of the time across all30 replicates within all pairwise combinations of the resource associated with the simplest function and the resources associated with the five next simplest functions. The probability of evolving the optimal phenotype at the intersection within a given resource pair varied from a low of 30% to a high of 100%, suggesting that it is easier for organisms to evolve some pairs of functions than others.

This outcome is likely the result of one of the valuable functions serving as a building block for the other function [15]. Such interaction effects illustrate the complex evolutionary dynamics that spatially heterogeneous environments can cause.

In our second round of experiments, we chose a single pair of resources, but tested all possible amounts of overlap. If resource patches have sufficiently small overlap, spatial niches should not be able to maintain a phenotypes capable of both functions. While phenotypes can extend beyond their target niche, their fitness will decrease in neighboring niches. As spatial niche size decreases, the probability that the matching phenotype will be lost from the population due to drift increases. As expected, the probability of the two-resource niche being occupied by organisms that could perform both valuable functions decreased with niche size. The configuration with 21 spaces between the centers of the circles, translating to a niche size of 50 cells, was the smallest at which this probability was greater than 50% (Fig. 2).

**Figure 2.**
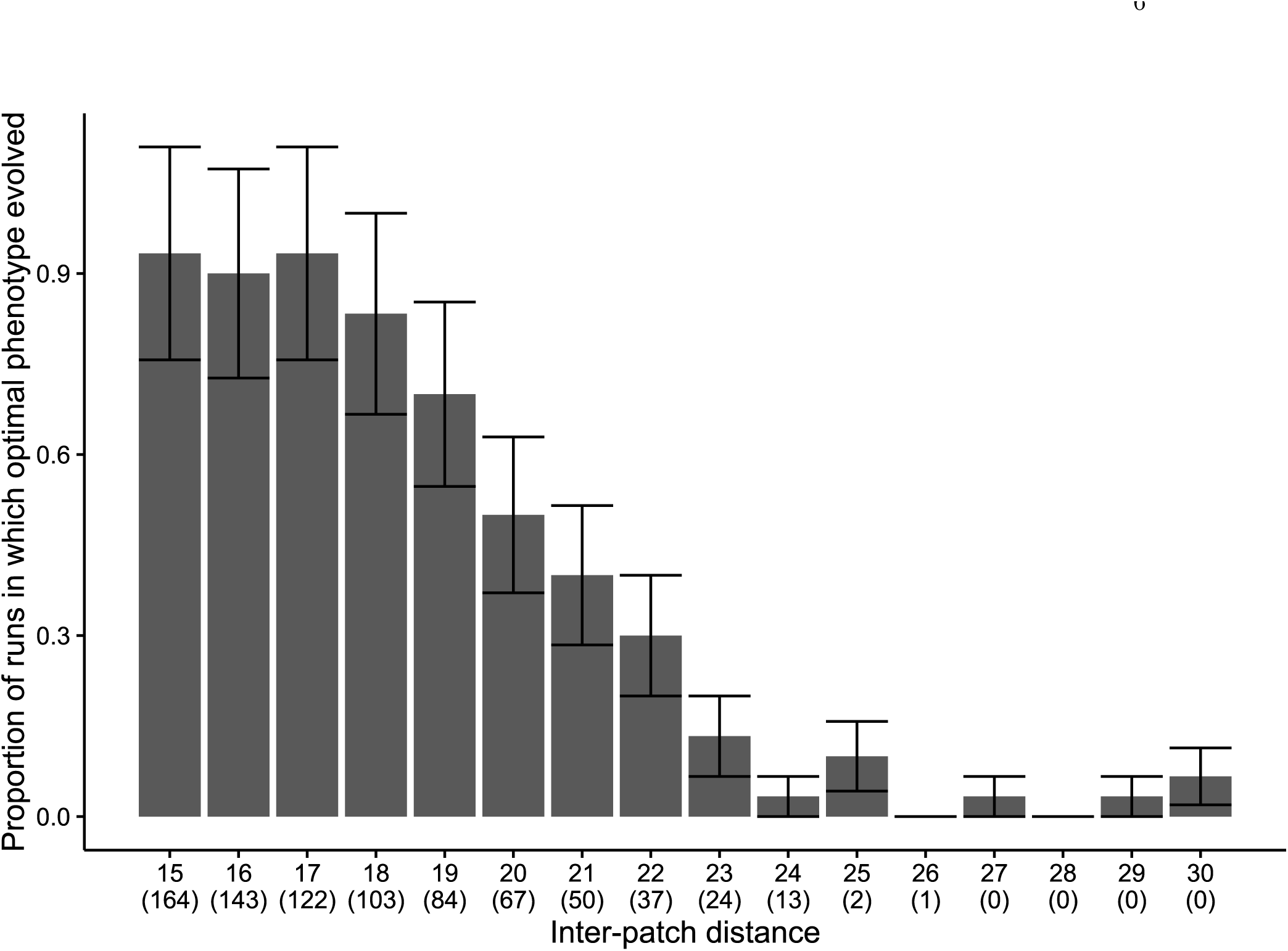
Proportion of populations (out of 30) at each inter-patch distance that contained organisms performing both functions (NOT and ORN) after 200,000 updates (approximately 4,000 generations). Error bars represent estimates of the standard deviation under the assumption that the underlying pattern resembles a Poisson distribution. Number of cells in the area where the patches overlap is specified in parenthesis for each distance.

### Effect of spatial heterogeneity on phenotypic diversity

Having established that sufficiently large overlaps between patches create new niches, we can now move on to more complex environments with more patches and more heterogeneity. We used three different techniques to arrange patches in the environment. In the first, we anchored patches in fixed positions and controlled spatial heterogeneity by adjusting patch size (Fig. 3). In the second, we gave all patches the same radius and controlled heterogeneity by setting the distance between their center points (Fig. 4).

**Figure 3.**
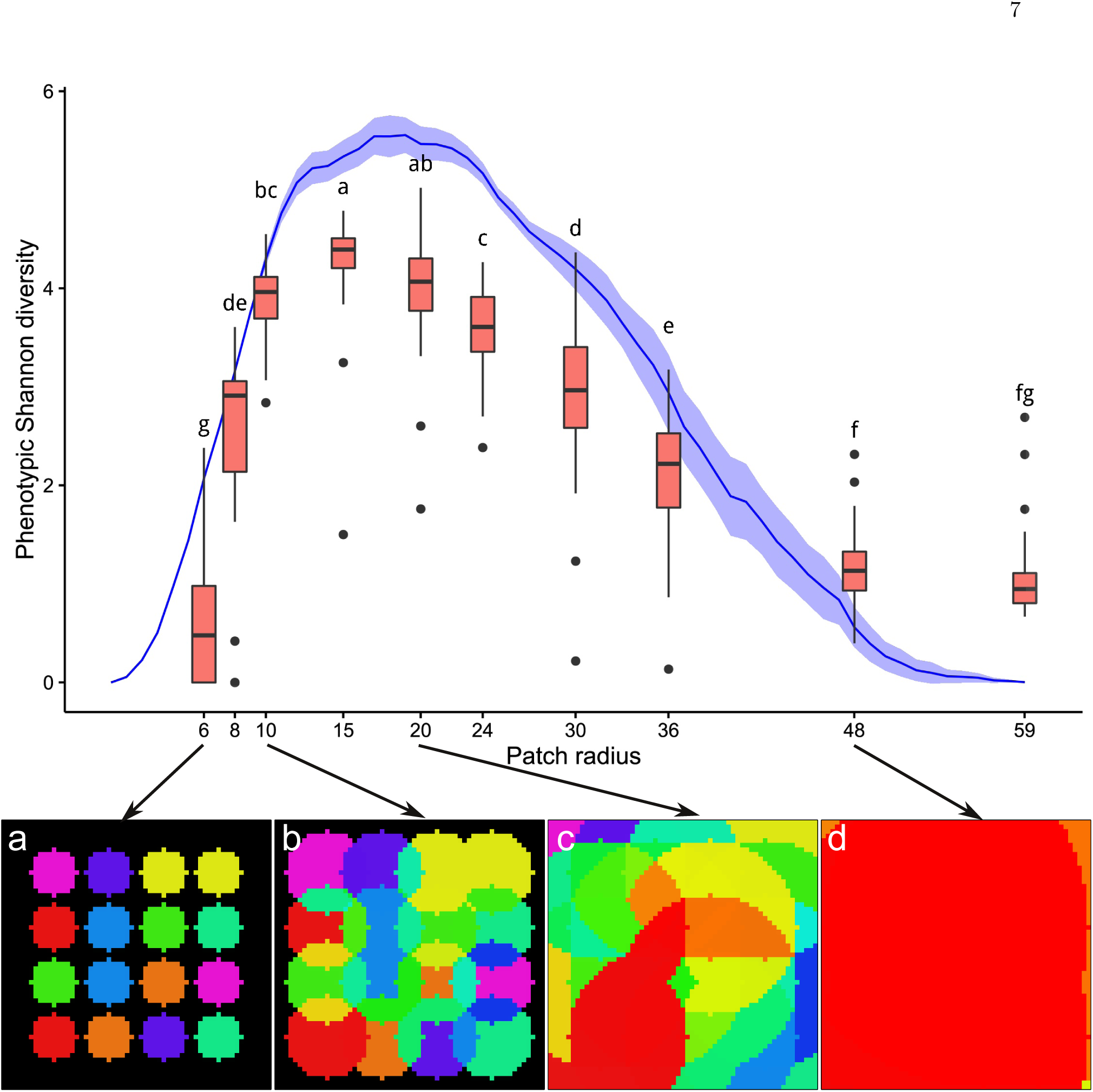
Shannon entropy (representing both phenotypic Shannon diversity and environmental entropy) across resource patch radii. The blue line shows environmental entropy at each radius (mean and standard deviation, calculated from 100 randomly generated environments at each radius). Environmental entropy peaks at patch radius 18. In environments with patch radii less than this peak (see sub-figures a and b), resources are more spatially isolated; conversely, in environments with patch radii greater than this peak, resources increasingly overlap (see sub-figure c). The high-radius extreme represents a homogeneous control environment (see sub-figure d), as all patches overlap and completely fill the world. Final phenotypic diversity at each radius condition tested in Avida is shown by the boxplots (n=30 for each radius; boxes show 25%, 50%, and 75% quantiles, with whiskers extending up to 1.5*IQR). Radii that do not share a letter have significantly different distributions of phenotypic diversity (pairwise two-sided Wilcoxon tests with Bonferroni adjustment, p<0.05). Sub-figures a-d show representative environments for the radius sizes indicated. Phenotypic diversity appears to peak at a patch radius somewhere between 13 and 23, an interval which includes the peak in environmental entropy.

In the third, we randomly selected the number, size, and location of patches in the environment, resulting in a variety of spatial heterogeneity levels (Fig. 5) (for details, see methods). The resource associated with the most complex logic function, EQU, was consistently available. We tracked the evolution of EQU as a proxy for the evolutionary potential of the population. However, the evolution of EQU should diminish the importance of spatial heterogeneity. To account for this confounding factor, we measured the diversity for each replicate as the phenotypic diversity of viable organisms in the population at the last time point prior to the evolution of EQU. For replicates in which EQU never evolved, we used phenotypic diversity from the final time point. To ensure that the presence of these two types of replicates did not bias the results, we repeated all analyses using data from the final time point for all replicates, and did not find any substantive difference.

Average phenotypic Shannon diversity differed significantly across resource patch sizes and proximities (Size: Kruskal-Wallis test, Chi-squared = 237.05, p< 2.2E-16, Fig. 3; Distance: Kruskal-Wallis test, Chi-squared = 104.0322, p *<* 2.2E-16, Fig. 4). Both size and distance experiments showed trends in phenotypic diversity qualitatively similar to the trends in spatial heterogeneity. The locations of the peaks in diversity overlapped with the peaks in environmental entropy (see Fig. 3 and Fig. 4).

**Figure 4.**
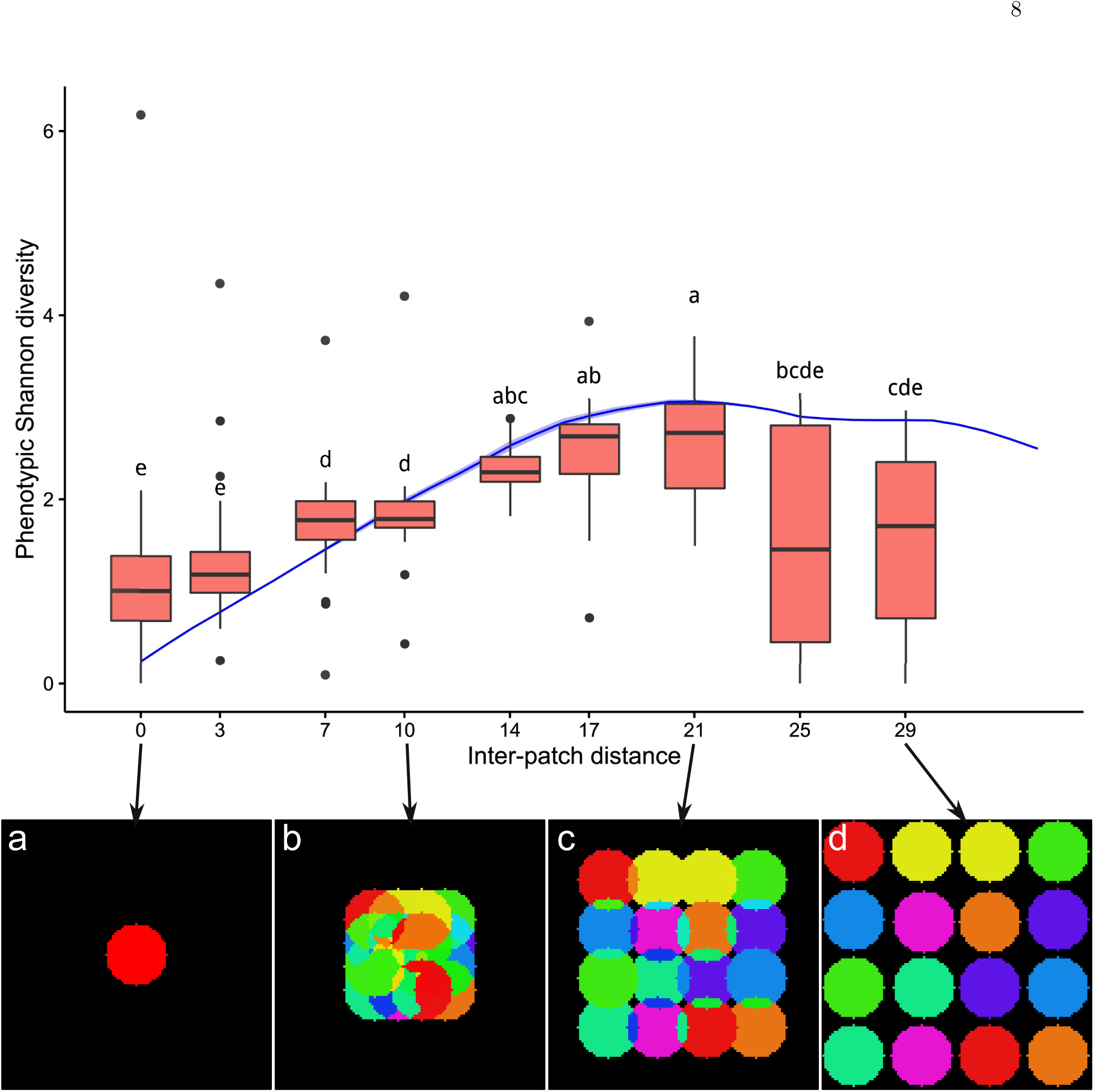
Shannon entropy (representing both phenotypic Shannon diversity and environmental entropy) across different resource inter-patch distances. The blue line shows environmental entropy at each distance (mean and standard deviation, calculated from 100 randomly generated environments at each distance; note that the standard deviation is small and hard to see). Environmental entropy peaks at inter-patch distance 21 (sub-figure c). In environments with inter-patch distances greater than this peak, resources are more spatially isolated (sub-figure d); conversely, at inter-patch distances lower than this peak, resources increasingly overlap (sub-figures a and b). Distance 0 represents a homogeneous control environment, as all resources overlap completely. Boxplots show final phenotypic diversity at each level tested in Avida (n=30 for each treatment; boxes show 25%, 50%, and 75% quantiles, with whiskers extending up to 1.5*IQR). Distances that do not share a letter have significantly different distributions of phenotypic diversity (pairwise two-sided Wilcoxon tests with Bonferroni adjustment, p<0.05). Sub-figures a-d show representative environments for the inter-patch distances indicated. Phenotypic diversity appears to peak at an inter-patch distance somewhere between 14 and 21, an interval which includes the peak in environmental entropy.

**Figure 5.**
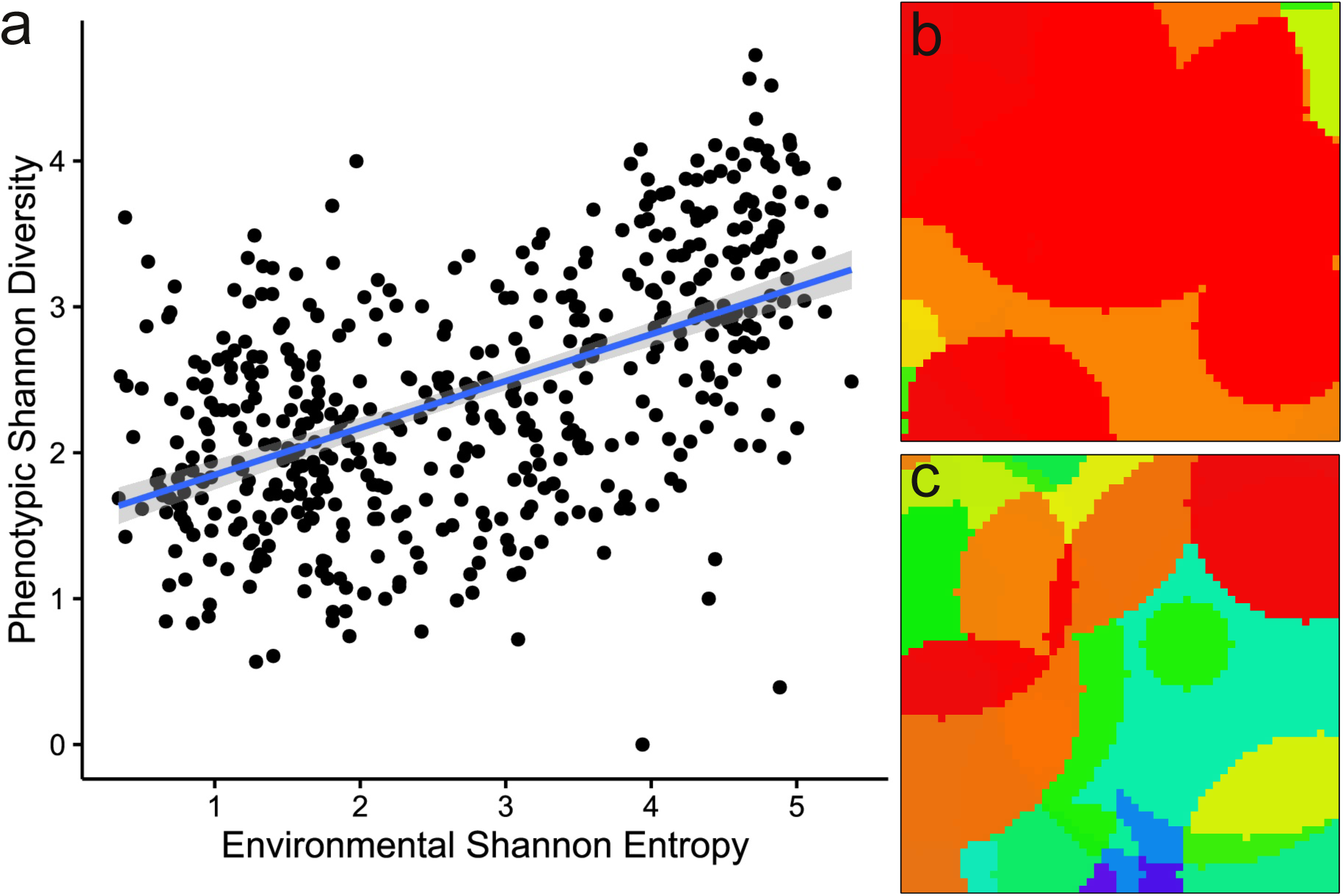
**A)** Scatterplot of environmental entropy vs phenotypic Shannon diversity across different randomly generated environments (Method 3) (n=500). Line represents the linear model shown in Table 1, and shaded area represents 95% confidence interval for the slope. **B)** and **C)** are two examples of randomly generated environments.

The third experimental treatment involved continuous rather than categorical variation in spatial heterogeneity, and we used a simple linear model regressing phenotypic Shannon diversity on environmental entropy to analyze the results (Table 1, 5). We also generated models including patch overlap as a metric of spatial configuration, but found no support for retaining the additional terms. Based on this model, we estimate that phenotypic diversity will increase by approximately one third of a unit, on average, for every unit that spatial heterogeneity increases - a fairly substantial effect.

**Table 1.** Regression table for model predicting phenotypic diversity, measured as Shannon entropy of viable organisms on the last measured update before EQU evolved (or the last update if EQU never evolved), from spatial heterogeneity, as measured by Shannon entropy of niches in the environment.

| y = Phenotypic Diversity |  |  |  |
| --- | --- | --- | --- |
| Variable | Estimate | SE | CI |
| Intercept | 1.53 *** | 0.07 | (1.39, 1.67) |
| Environmental Entropy | 0.32 *** | 0.02 | (0.28, 0.37) |

Residual standard error: 0.71 on 498 DF $R^2 = 0.28$ , F-statistic = 196.3 on 1 and 498 DF

Across the board, populations that evolved in spatially heterogeneous environments were more diverse than those evolved in more homogeneous environments, regardless of how that heterogeneity was generated. Increasing spatial heterogeneity partitions resources into spatially structured areas, thereby creating new, unique niches into which organisms can diversify and partially escape competition. This trend is in contrast to the peak in diversity found at intermediate heterogeneity by some other experiments [12,13]. We suspect that this result occurred because, in our experiments, increased heterogeneity does not necessarily come at the cost of decreased fundamental niche size. Our results agree with the many biological studies that have found a positive relationship between phenotypic diversity and spatial heterogeneity [3].

### Spatial heterogeneity and evolutionary potential

The proportion of replicates in which the most complex task (EQU) evolved had a positive saturating response to increasing patch radius. As patch radius increased from low (6-10) to intermediate values (15-24), populations evolved the EQU function significantly more frequently. (Fig. 6). Evolutionary potential reached a maximum saturating response such that we were not able to detect a significant difference between the probabilities of EQU evolving in trials with medium and high patch radii *(>* 24) after a Bonferroni correction (Fig. 6).

**Figure 6.**
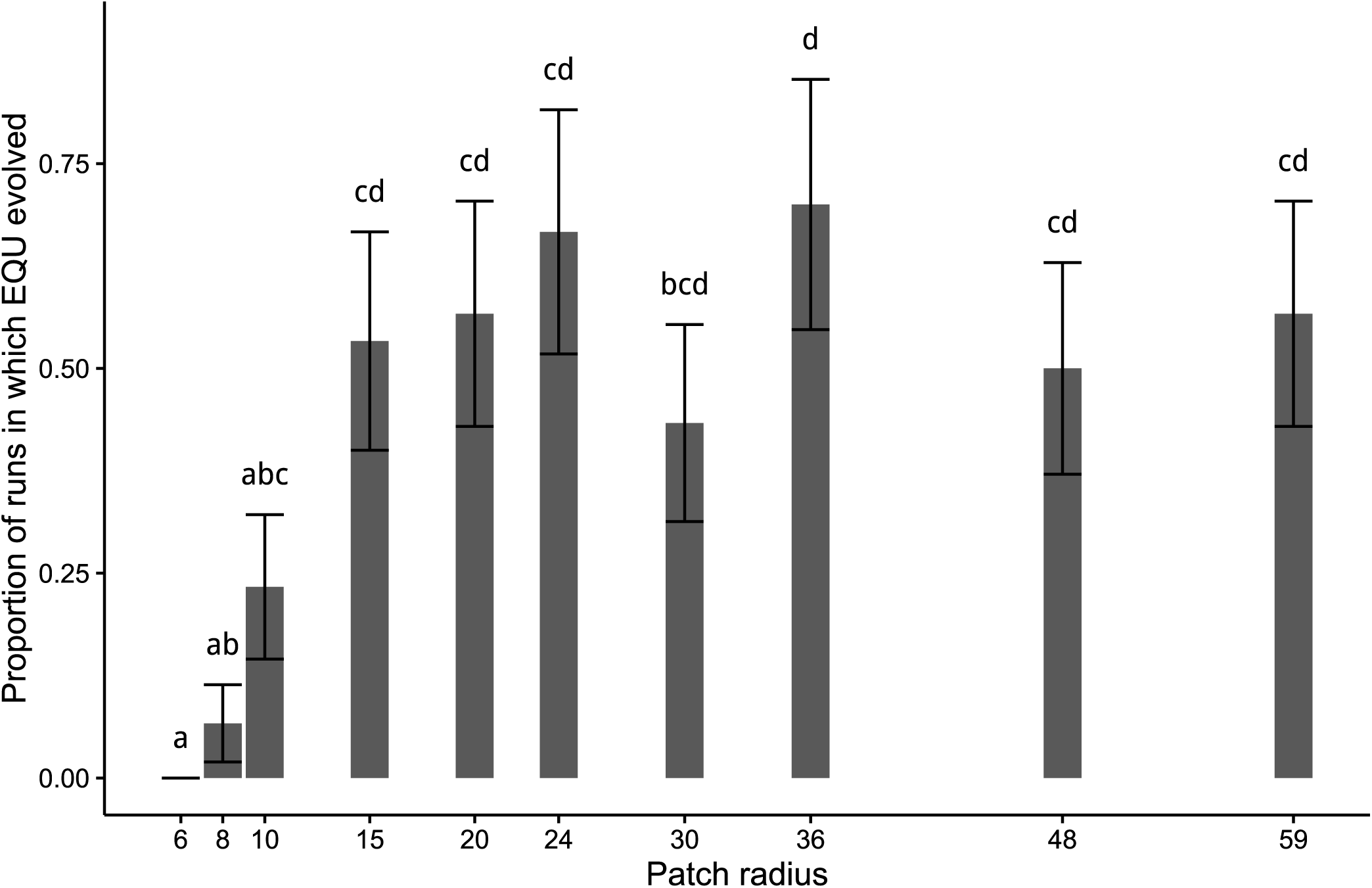
Proportion of populations (n=30 per treatment) that evolved EQU across different patch radius sizes. Error bars are estimates of standard deviation based on the square root of the count of populations evolving EQU for each condition. Radii that do not share a letter have significantly different probabilities of evolving EQU (pairwise two-sided Fisher’s exact tests with Bonferroni adjustment, p<0.05).

The inter-patch distance experiment highlighted the importance of patch overlap relative to spatial entropy in evolving EQU. At low distances between patch centers (0-14), resource patch overlaps and evolutionary potential were at their highest levels (Fig. 7). At intermediate to large inter-patch distances, evolutionary potential decreased, as did the intersection between patches. Larger overlaps have multiple resources available, allowing populations to build up to more complicated functions with greater ease than than in conditions in which patches overlap less [15].

**Figure 7.**
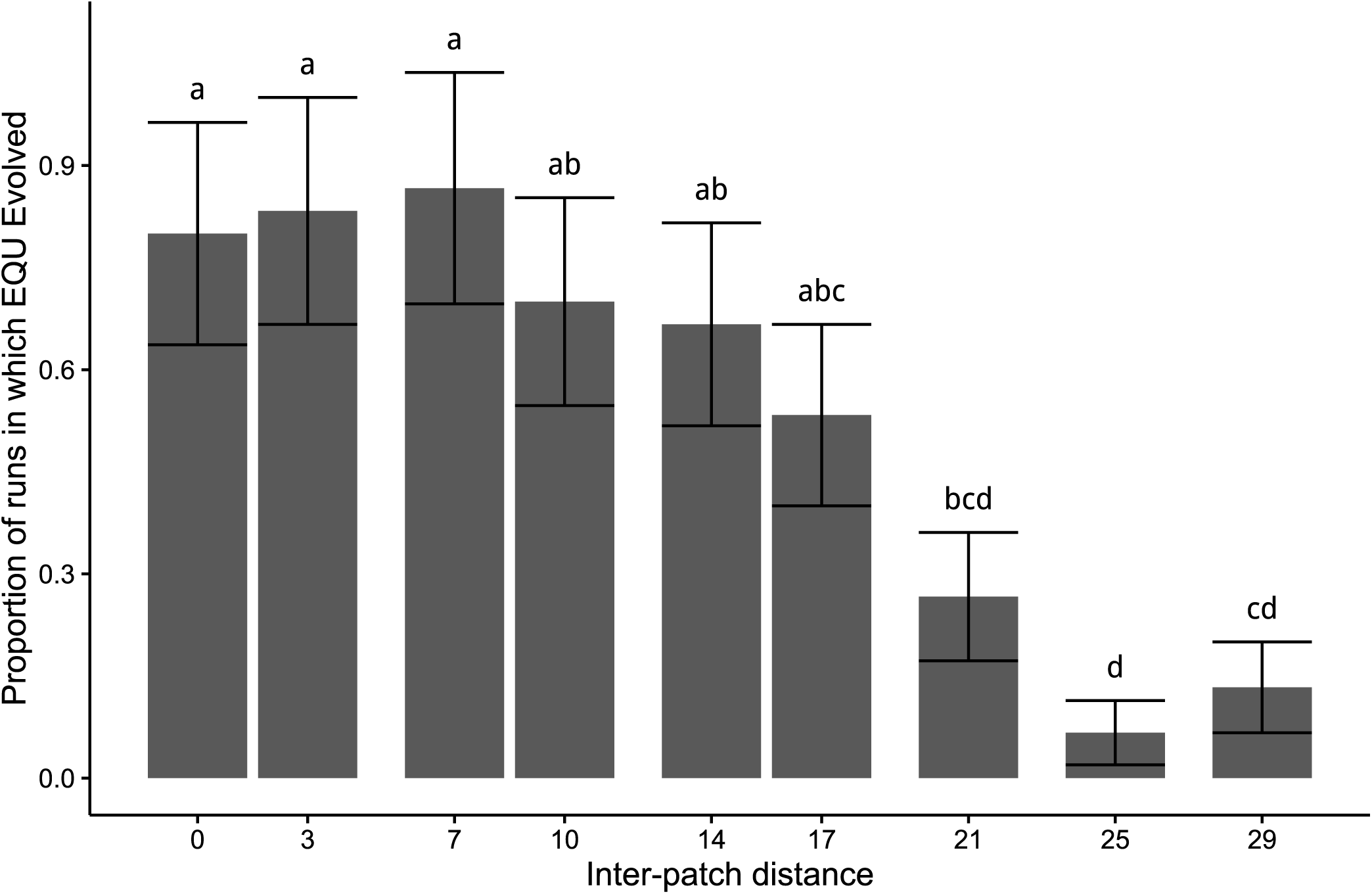
Proportion of populations (n=30 per treatment) that evolved EQU across different inter-patch distances. Error bars are estimates of standard deviation based on the square root of the count of populations evolving EQU for each condition. Inter-patch distances that do not share a letter have significantly different probabilities of evolving EQU (pairwise two-sided Fisher’s exact tests with Bonferroni adjustment, p<0.05).

The common trend from the first two experiments is that the amount of overlap between resource patches (i.e. the number of different resources present within a spatial niche) has a stronger impact on evolutionary potential than diversity. To disentangle the effect of resource overlap (our spatial configuration metric) and spatial heterogeneity (our spatial composition metric) on evolutionary potential, we built a linear model using the overlap as a predictor. Recall from our discussion of Quantifying Heterogeneity that we measured overlap by looking at the distribution of the number of resource types per cell in an environment, and taking its mean, variance, skew, and kurtosis. Ultimately, these statistics capture similar information about the environment to that captured by spatial entropy, as demonstrated by the high *R^2^* value of the linear model predicting environmental entropy from the mean and skew of number of resources available per cell (see Table 2). Variance and kurtosis were dropped from the final model because of the high colinearity between mean and variance and between skew and kurtosis. Thus, we did not include spatial entropy in the overall model of evolutionary potential, as it would have added largely redundant information.

**Table 2.**
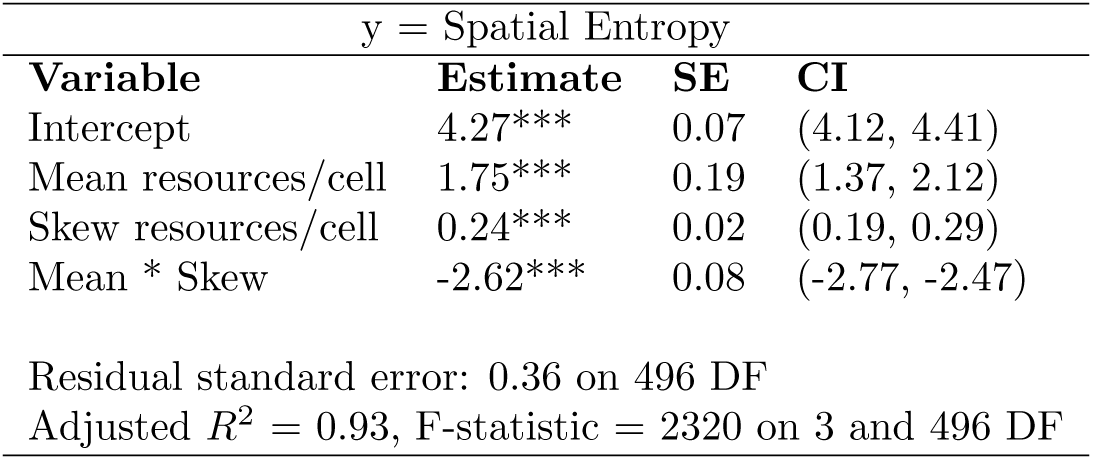
Regression table for linear model predicting entropy of the environment from the mean and skew (and their interaction) of the number of resource types per cell.

We used a logistic regression to predict the probability of evolving the EQU function given phenotypic diversity and environmental heterogeneity (Table 3). As environmental descriptors, we used mean number of resources per cell, skew of the number of resources per cell, and their interaction. We included phenotypic diversity (before EQU evolved) as the final predictor in the model of evolutionary potential, due to its positive association with spatial heterogeneity. As expected, this model suggests that diversity is important to evolutionary potential. Every additional unit of phenotypic diversity increases the odds of EQU evolving by approximately 4.3 times. However, this effect is dwarfed by the effect of mean resources per cell (i.e. overlap). Increasing the average number of resources per cell by 1 (e.g. adding an additional resource patch that covers the entire grid) would increase the odds of EQU evolving by approximately 148.8 times. The interaction between diversity and mean resources per cell has an effect of a similar magnitude. This interaction suggests that either A) environments with more overlap amplify the benefits of diversity to evolutionary potential, B) environments with more overlap promote the evolution of diversity that is more useful to evolving EQU than the diversity promoted by other environments, or C) both.

**Table 3.** Logistic regression table for linear model predicting whether or not EQU evolved from phenotypic diversity, mean resource types per cell, and skew of resource types per cell, as well as all interactions between these variables.

| y = EQU Evolution |  |  |  |
| --- | --- | --- | --- |
| Variable | Estimate | SE | CI |
| Intercept | 1.30*** | 0.17 | (0.97, 1.66) |
| Phenotypic Diversity | 1.46*** | 0.22 | (1.05, 1.90) |
| Mean resources/cell | 5.00*** | 0.84 | (3.40, 6.71) |
| Skew resources/cell | 0.74*** | 0.21 | (0.32, 1.13) |
| Diversity * Mean | 4.38*** | 0.93 | (2.60, 6.25) |
| Diversity * Skew | 0.81** | 0.26 | (0.30, 1.32) |
| Mean * Skew | 1.70 | 0.81 | (-0.00, 3.21) |
| Diversity * Mean * Skew | 2.65** | 1.01 | (0.64, 4.62) |
| Null deviance: 651.08, 499 DF, Residual deviance: 549.88, 492 DF |  |  |  |

Ultimately, the amount of overlap between single-resource patches is the strongest driver of the evolutionary potential of EQU, with global phenotypic diversity playing an important secondary role. This impact of diversity may be responsible for the saturating evolutionary potential curves seen in the first two experiments; during the plateaus in these curves, overlap is decreasing while diversity is increasing. Given the strength of the synergies between functions in Avida, it is possible that the effect of overlap in this system is greater than it would be in other contexts. These results support the idea that it is possible to select for diversity that is more or less useful to evolving a specific feature.

## Conclusion

We have presented evidence supporting the positive diversity-heterogeneity hypothesis. We found that population diversity increases in response to higher levels of spatial heterogeneity, rather than peaking at an intermediate level. Further, we have shown that this increase in diversity leads to higher evolutionary potential of a complex function. However, the spatial configuration of the landscape in which a population is evolving can have a much stronger effect on evolutionary potential than diversity alone. The presence of a synergistic interaction between diversity and spatial configuration provides preliminary evidence that landscape configuration can promote diversity that is more or less useful to evolving a given trait. These findings help to solidify the growing consensus that there is a causal relationship between spatial heterogeneity and population diversity. Moreover, by exploring these changes in diversity in an evolutionary context, we push toward a more mechanistic understanding of spatial heterogeneity and diversity.

However, a number of open questions remain. Most notably, we only considered diversity as phenotypic Shannon diversity within individual populations. It is possible that other measures of diversity, such as beta-diversity across populations, would have different relationships to spatial heterogeneity. Similarly, the only metric of spatial heterogeneity that we explored was Shannon entropy. There are many alternative metrics, particularly metrics of spatial composition (see Quantifying Heterogeneity), that may have interesting subtleties in their relationship to diversity. Additionally, allowing sexual recombination, motility, or the evolution of dispersal distance could produce substantially different results.

Our results have implications beyond ecology; for example these same dynamics can be harnessed to improve computational problem solving with evolutionary algorithms. A common approach to solving hard problems in these systems is to first evolve solutions to easier related problems and use them as building blocks [15,30,31]. Spatially heterogeneous environments can be created by rewarding population members that solve a range of simpler problems in different parts of a spatially structured population. Recent research on an evolutionary algorithm that creates spatial heterogeneity via host-parasite co-evolution supports the idea that this approach may be useful [32]. Thus, it may be possible to use the results of our study to create environments that are particularly conducive to solving problems with evolutionary computation. Such an approach would synergize well with previously discovered positive impacts of spatial structure on diversity maintenance in evolutionary computation [33,34].

Understanding drivers of diversity, particularly with regard to their impact on evolutionary potential, also has important practical implications for preserving biodiversity in natural systems. Landscapes world-wide are experiencing new disturbance regimes and there is currently little theory predicting how ecosystems will respond [35, 36]. Anthropogenic disturbances tend to be a homogenizing force [37], but this effect can be mitigated by urban planning if we understand the evolutionary and ecological dynamics at play. Given the theorized importance of biodiversity to ecosystem stability and function, knowing how spatial heterogeneity can affect diversity across different ecosystems will be critical to achieving such goals [38, 39]. Thus, going forward, it will be important to solidify our understanding of the factors affecting the shape of the relationship between spatial heterogeneity and diversity.

## Methods

These experiments were carried out using Avida 2.12.4, an open-source platform for digital evolution [14]. Digital organisms in Avida have genomes made up of simple computer instructions. These instructions comprise a language that is computationally universal; there are no fundamental limits to the types of algorithms they can implement. Each organism runs in a virtual computer processor. The world, a grid of cells that can each be occupied by a single organism, is seeded with organisms that can do nothing except replicate themselves. When instructions are copied during this replication process, they may be stochastic ally mutated. These mutations are a source of variation, allowing organisms to evolve to replicate faster or use energy (CPU cycles) more efficiently as they execute their genomes. They can get more CPU cycles by metabolizing resources; these metabolic activities involve performing computational functions (Boolean logic) specific to the resource type. Some of the metabolic functions are more challenging to evolve than others, based on the complexity of the code required to perform the associated function.

In our experiments, we have nine geographically-limited resources on a 59x59 grid. Organisms that perform a logic function must be in an area with the associated resource to successfully metabolize it and receive the additional CPU cycles. In this study, we controlled spatial heterogeneity by varying the geographic limitations associated with each type of resource. We placed circular patches of single resources in the environment, some of which overlapped to create new multi-resource spatial niches. To maintain a consistent measure for evolutionary potential, we placed the resource associated with the most complex logic function everywhere.

Specific details for experimental treatments are described below. For each, we performed 30 replicates and initialized all configuration settings to Avida defaults (unless otherwise specified) with the exception of copy mutations, which we set to a probability of 0.0025 per instruction copied. We allowed each replicate to evolve for 100,000 updates. An update is the amount of time it take for the average organism in the population to execute 30 instructions; 100,000 updates translates to approximately 30,000 generations on average.

We recorded phenotypic Shannon diversity of organisms and the number of populations that had at least one organism that performed the most complex logic function at the end of the experiment. We define an organisms phenotype as the set of logic functions it performs. We performed all statistics in R version 3.0.1 [40,41]. For each experimental condition, we ran a large number of replicates relative to the amount of variation in results, ensuring sufficient power to detect reasonably-sized effects. Because each replicate had a unique random number generator seed, all data points were independent and, as such, the assumptions for all statistical tests were met.

### Niche stability

For the niche stability experiments, each circle had a radius of 14 cells. We set up the patches to be perfectly on top of each other in the complete-overlap condition (0 cell distance between centers), 14 cells between centers for halfway overlap, and 30 cells for no overlap. We tested these conditions in a full-factorial design with 30 replicates of all potential pairwise combinations of resources, for a total of 192 treatments.

We used the two simplest logic functions in the functional niche-size assessment experiments because both functions consistently evolved in all of the two-circle overlap experiments where they were present. These patches also had a radius of 14, and between 0 and 30 cells between their centers. To ensure that niches created in this experiment were ecologically stable, we allowed these replicates to continue for an additional 100,000 update”ecology phase” with no mutations. We measured niche stability as the frequency with which intermediate phenotypes evolved and persisted until the end of the experiment.

### Fixed layout experiments

To systematically measure the effect of spatial heterogeneity on diversity, we ran two sets of experiments with 16 resource patches (two for each resource type) placed at pre-determined locations in the environment. In the *patch radius experiment*, we varied the size of each circular patch, while in the *inter-patch distance experiment*, we varied the spacing between fixed-sized patches. To select treatment values for each of these variables, we wrote a simulation in Python that created random environments with the specified parameters and measured their heterogeneity (available at <u>https://github.com/emilydolson/resourceheterogeneity)</u>. The results of these runs produced the data for the curves in figures 3 and 4. We used these curves to select values of each variable that would produce a range of levels of environmental heterogeneity. Across both sets of experiments, the most complex logic function was valuable everywhere to ensure that it would be maintained in the population long enough to be measured if it evolved. We designed many treatments within each experiment (detailed below) and ran 30 replicates for each treatment. While the positions of each resource patch were consistent within a treatment, we randomized the type of resource in each patch across replicates.

In the patch-radius experiment, we manipulated spatial resource heterogeneity by changing the radius sizes of the single-resource patches. We placed patch centers 12 spaces apart. Based on the results of the Python simulation, we used patch radii of 6, 8, 10, 15, 20, 24, 30, 36, 48, and 59 as our experimental conditions (see Fig 3). Changing patch radii introduces a potentially confounding variable: the overall amount of resources in the environment. To control for this additional factor, we ran another experiment in which we manipulated spatial resource heterogeneity by changing the distances between the resource patches while holding patch radius constant. At each inter-patch distance, we set resource patches to a constant size (radius 14) in a 115 x 115 non-toroidal world. We manipulated distances between patch centers across treatments (using distances 0, 3, 7, 10, 14, 17, 21, 24, or 29) (see Fig 4). This condition had a constant quantity of each resource, but introduced its own constraints on the types of environments that could exist at each level of heterogeneity. In combination with each other, however, these two sets of experiments address the effect of heterogeneity in a more comprehensive way.

We used Kruskal-Wallis tests to determine if the distribution of phenotypic Shannon diversity varied significantly across treatments. Subsequently, we determined where average diversity peaked using post-hoc pairwise Wilcoxon tests with a Bonferroni correction for multiple comparisons. We used Fishers exact tests to determine if the number of populations that performed the the most complex logic function by the end of the experiment changed across distance treatments. We performed additional Fishers exact tests to determine where evolutionary potential peaked.

### Randomized patch experiments

In the two prior experiments, it is hard to disentangle the specific effects of patterned resource overlap from the more general effects of heterogeneity, as these variables covaried. To sort out these factors, we placed a random number of patches of random sizes in random locations in the environment. Using the Python simulation from the previous section to determine parameter ranges that would produce the widest variety of levels of spatial heterogeneity, we tested 500 environmental configurations, consisting of 4-124 patches with radii between 6 and 35. Ultimately, the total number of resource patches in an environment was the primary determiner of its heterogeneity; varying patch sizes and placements achieved variation in patch overlap within a given level of heterogeneity.

Because these experiments produced continuous rather than categorical data, we used linear regression models in R to determine the factors influencing phenotypic diversity and spatial heterogeneity. For the logistic regression predicting evolution of EQU, we used a generalized linear model. We narrowed down potential models using AIC; we then evaluated and trimmed factors with high collinearity until we found the minimally adequate model. We also fit a model that included total sum of resources across all cells, but there was a curvilinear relationship between total sum of resources and spatial entropy that rendered the factors largely redundant and added little explanatory power. We identified a substantial outlier, in the form of a single run in which the population went extinct, but removing it had negligible effect on our statistical analyses, so it was included.

### Code Availability

Source code for Avida is available at available at <u>http://avida.devosoft.org</u>. These experiments were carried out using Avida version 2.12.4. Scripts used to generate different environmental conditions and analyze data are available via Zenodo with DOI [removed for double-blind submission]

## Data Availability

The data supporting the findings of this study are available via Zenodo with DOI [removed for double-blind submission]

## References

1. Hutchinson GE (1961) The paradox of the plankton. American Naturalist: 137–145.

2. Chesson P (2000) Mechanisms of maintenance of species diversity. Annual Review of Ecology and Systematics 31: 343–366.

3. Stein A, Gerstner K, Kreft H (2014) Environmental heterogeneity as a universal driver of species richness across taxa, biomes and spatial scales. Ecology letters 17: 866–880.

4. Pianka ER (1966) Convexity, Desert Lizards, and Spatial Heterogeneity. Ecology 47: 1055–1059.

5. Pianka ER (1966) Latitudinal Gradients in Species Diversity: A Review of Concepts. The American Naturalist 100: 33–46.

6. Pacala S, Tilman D (1994) Limiting similarity in mechanistic and spatial models of plant competition in heterogeneous environments. American Naturalist 143: 222–257.

7. Jankowski K, Schindler DE, Horner-Devine MC (2014) Resource availability and spatial heterogeneity control bacterial community response to nutrient enrichment in lakes. PLoS ONE 9: e86991.

8. Barnett A, Beisner BE (2007) Zooplankton biodiversity and lake trophic state: explanations invoking resource abundance and distribution. Ecology 88: 1675–1686.

9. Tilman D (1977) Resource competition between plankton algae: An experimental and theoretical approach. Ecology 58: 338–348.

10. Harpole WS, Tilman D (2007) Grassland species loss resulting from reduced niche dimension. Nature 446: 791–793.

11. Allouche O, Kadmon R A general framework for neutral models of community dynamics 12: 1287–1297.

12. Allouche O, Kalyuzhny M, Moreno-Rueda G, Pizarro M, Kadmon R Areaheterogeneity tradeoff and the diversity of ecological communities 109: 17495–17500.

13. Haller BC, Mazzucco R, Dieckmann U Evolutionary branching in complex landscapes. The American Naturalist 182: E127–E141.

14. Ofria C, Wilke CO (2004) Avida: A software platform for research in computational evolutionary biology. Artificial Life 10: 191–229.

15. Lenski RE, Ofria C, Pennock RT, Adami C (2003) The evolutionary origin of complex features. Nature 423: 139–144.

16. Wilke CO, Wang JL, Ofria C, Lenski RE, Adami C (2001) Evolution of digital organisms at high mutation rates leads to survival of the flattest. Nature 412: 331–333.

17. Chow SS, Wilke CO, Ofria C, Lenski RE, Adami C (2004) Adaptive radiation from resource competition in digital organisms. Science 305: 84–86.

18. Fortuna MA, Zaman L, Wagner AP, Ofria C (2013) Evolving digital ecological networks. PLoS Comput Biol 9: e1002928.

19. Cooper TF, Ofria C (2003) Evolution of stable ecosystems in populations of digital organisms. In: Artificial Life VIII: Proceedings of the Eighth International Conference on Artificial life. pp. 227–232.

20. Schluter D (2009) Evidence for ecological speciation and its alternative. Science 323: 737–741.

21. Lavergne S, Molofsky J (2007) Increased genetic variation and evolutionary potential drive the success of an invasive grass. Proceedings of the National Academy of Sciences 104: 3883–3888.

22. McDonald BA, Linde C (2002) Pathogen population genetics, evolutionary potential, and durable resistance. Annual Review of Phytopathology 40: 349–379.

23. De Jong KA (1975) Analysis of the behavior of a class of genetic adaptive systems.

24. Goldberg DE, Richardson J Genetic algorithms with sharing for multimodal function optimization. In: Genetic algorithms and their applications: Proceedings of the Second International Conference on Genetic Algorithms. Hillsdale, NJ: Lawrence Erlbaum, pp. 41–49.

25. Lehman J, Stanley KO (2008) Exploiting open-endedness to solve problems through the search for novelty. In: Proceedings of the Eleventh International Conference on Artificial Life (ALIFE XI). Cambridge, MA: MIT Press.

26. Mouret JB, Doncieux S Using behavioral exploration objectives to solve deceptive problems in neuro-evolution. In: Proceedings of the 11th Annual Conference on Genetic and Evolutionary Computation. ACM, GECCO ‘09, pp. 627–634. doi: 10.1145/1569901.1569988. URL http://doi.acm.org/10.1145/1569901.1569988.

27. Walker BL, Ofria C (2012) Evolutionary potential is maximized at intermediate diversity levels. MIT Press, pp. 116–120. doi:10.7551/978-0-262-31050-5-ch017.

28. McGarigal K (2006) Landscape pattern metrics. In: Encyclopedia of Environmetrics, John Wiley & Sons, Ltd.

29. Forman RTT, Godron M (1986) Landscape ecology. Wiley.

30. Bongard JC, Hornby GS (2010) Guarding Against Premature Convergence While Accelerating Evolutionary Search. In: Proceedings of the 12th Annual Conference on Genetic and Evolutionary Computation. New York, NY, USA: ACM, GECCO ‘10, pp. 111–118. doi:10.1145/1830483.1830504. URL <u>http://doi.acm.org/10.1145/1830483.1830504.</u>

31. Grabowski L, Magaa J (2014) Building on Simplicity: Multi-stage Evolution of Digital Organisms. The MIT Press, pp. 113–120. doi:10.7551/978-0-262-32621-6-ch019.

32. Harper R (2012) Spatial co-evolution: Quicker, fitter and less bloated. In: Proceedings of the 14th Annual Conference on Genetic and Evolutionary Computation. ACM, GECCO ‘12, pp. 759–766. doi:10.1145 / 2330163.2330269.

33. Whitley D, Rana S, Heckendorn RB (1998) The island model genetic algorithm: On separability, population size and convergence. Journal of Computing and Information Technology 7: 3347.

34. Cant-Paz E, Goldberg DE (2003) Are multiple runs of genetic algorithms better than one? In: Cant-Paz E, Foster JA, Deb K, Davis LD, Roy R, et al. editors, Genetic and Evolutionary Computation GECCO 2003, Springer Berlin Heidelberg, number 2723 in Lecture Notes in Computer Science. pp. 801–812.

35. Arnell NW (2004) Climate change and global water resources: SRES emissions and socio-economic scenarios. Global Environmental Change 14: 31–52.

36. Galloway JN, Dentener FJ, Capone DG, Boyer EW, Howarth RW, et al. (2004) Nitrogen cycles: past, present, and future. Biogeochemistry 70: 153–226.

37. McKinney ML (2006) Urbanization as a major cause of biotic homogenization. Biological Conservation 127: 247–260.

38. Craine JM, Ocheltree TW, Nippert JB, Towne EG, Skibbe AM, et al. (2013) Global diversity of drought tolerance and grassland climate-change resilience. Nature Climate Change 3: 63–67.

39. Wilmers CC, Getz WM (2005) Gray wolves as climate change buffers in yellowstone. PLoS Biology 3: e92.

40. Wickham H (2009) ggplot2: elegant graphics for data analysis. Springer New York. URL http://had.co.nz/ggplot2/book.

41. R Core Team (2014) R: A Language and Environment for Statistical Computing. R Foundation for Statistical Computing, Vienna, Austria. URL http://www.R-project.org.

